# Do silver-chitosan nanocomposites promote bacterial resistance to silver or common antibiotics?

**DOI:** 10.1101/2024.06.13.596383

**Authors:** Mariliis Sihtmäe, Jüri Laanoja, Maarja Otsus, Heiki Vija, Anne Kahru, Kaja Kasemets

**Affiliations:** Laboratory of Environmental Toxicology, National Institute of Chemical Physics and Biophysics, Akadeemia tee 23, 12618 Tallinn, Estonia; Department of Chemistry and Biotechnology, School of Science, Tallinn University of Technology, Ehitajate tee 5, 19086 Tallinn, Estonia; Estonian Academy of Sciences, Kohtu 6, 10130 Tallinn, Estonia

**Keywords:** hospital-acquired infections, chitosan, silver, nanocomposites, antibiotics, antimicrobial resistance development

## Abstract

Silver-containing nanomaterials are versatile antimicrobials for slowing down the rapid increase of nosocomial infections caused by antibiotic-resistant bacteria. However, silver can also promote resistance in bacteria to both silver itself and conventional antibiotics. Despite this, the topic is still poorly studied.

Silver-chitosan nanocomposites (nAgCSs) with a primary size of ∼50 nm and weight ratios of 1:1 (nAgCS-1) and 1:3 (nAgCS-3) were synthesised and studied for potential antimicrobials, such as wound dressings, antimicrobial coatings. These nAgCSs exhibited antibacterial efficacy at level of 6.25–14.6 mg Ag/L level (minimal inhibitory concentration, MIC) against *Escherichia coli* and *Staphylococcus aureus* being comparable to the efficiency of AgNO_3_ (MIC 3.13–7.29 mg Ag/L) or benzalkonium chloride (MIC 0.91–6.25 mg/L). Low molecular weight chitosan also demonstrated antibacterial activity against *S. aureus* (MIC 25 mg/L), though it was less effective against *E. coli* (MIC 70 mg/L). Notably, nAgCS-3 was almost as efficient as AgNO_3_ against *S. aureus* (MIC 6.25–7.29 mg/L) and benzalkonium chloride against *E. coli* (MIC 6. mg/L). Confocal laser scanning microscopy revealed a noticeable aggregation of *S. aureus* cells caused by exposure to nAgCS-3, an effect that was less pronounced in *E. coli*.

The development of bacterial resistance to nAgCSs and AgNO_3_ and cross-resistance to 14 conventional antibiotics upon continuous exposure (up to 5 weeks) to sub-inhibitory concentrations (EC_20_) of the studied silver compounds was examined. No resistance development was observed for the studied silver compounds and antibiotics in either *E. coli* or *S. aureus*. Thus, silver-chitosan nanocomposites show promise as efficient and not AMR-inducing compounds for antimicrobial applications, such as wound dressings and surfaces.

## 1. Introduction

Antimicrobial resistance (AMR) is an urgent global public health threat. According to the United Nations report, by 2050, the prognosed number of human deaths due to AMR will be comparable to that of cancer – both causing about 10 million deaths annually [1]. Moreover, during the COVID-19 pandemic, antibiotics were excessively prescribed to avoid secondary bacterial infections, possibly boosting the spread of AMR [2]. Furthermore, the clinical development of new antimicrobials is almost at a standstill. In 2019, the World Health Organization (WHO) identified 32 antibiotics under clinical development, addressing the WHO list of priority pathogens. Out of these 32, only six were classified as innovative. In addition, a lack of access to quality antimicrobials, especially in healthcare systems, also remains a major issue affecting countries of all levels of development [3]. Despite that, the global consumption of antibiotics in humans has risen in the past two decades.

Nanotechnology promises to create novel, efficient, and safe antimicrobials for biomedical applications, e.g., wound dressings and implants, that can reduce both infections and the development of antibiotic-resistant bacteria. Silver (Ag) compounds, including various forms of nanosilver, are widely used in wound dressings and disinfectants in health care due to their antibacterial activity and assumingly low risk of development of AMR [4,5].

Silver-based compounds have been used for their antiseptic properties for centuries [6]. Silver (incl. in nano form) is used for treating infections encountered in burns, open wounds, and chronic ulcers. It has been shown that the size, shape, surface coating and surface charge of silver nanomaterials determine their antimicrobial activity and efficacy [7–10]. It is also known that the toxicity of nanosilver is driven mainly by the dissolved ions, and bioavailability and toxicity increase when in contact with bacterial cells [11]. In the case of silver nanomaterials, the release of Ag ions can be controlled, reducing bacterial growth and the likelihood of resistance development [12].

Indeed, there are only a few reports of bacterial resistance to ionic silver, with one of the earliest reported by Jelenko *et al*. [13], where resistant *Escherichia coli* was isolated from a clinical case of prolonged AgNO_3_-treated burns. This finding was followed by the discovery of a silver-resistant *Salmonella typhimurium* strain in the 1970s, also isolated from AgNO_3_-treated burns [14]. *E. coli* and *S. aureus* are currently among the leading global causes of nosocomial infections caused by antibiotic-resistant bacteria [15,16].

In general, examples of bacteria adapting to conditions of prolonged exposure to silver are found in specific environmental settings where toxicity might select for resistance [17], including the soil of a silver mine, photographic laboratory effluent [18], practices related to dental amalgams, burn units where silver-containing skin disinfectants are applied, and the use of silver-coated catheters [19]. However, bacterial resistance to silver nanoparticles/nanomaterials has not been extensively studied, and the available data are controversial and inconclusive (see below).

Recently, a report by Panacek *et al*. [20] was published showing the development of silver resistance in three different Gram-negative bacteria (two *E. coli* and one *Pseudomonas aeruginosa* strain) after repeated exposure in 20 consequent culture steps to subinhibitory concentrations of silver nanoparticles (NPs). The developed resistance was phenotypic and was triggered by the production of flagellin – an extracellular bacterial protein forming motility-enabling flagellar filaments – that caused the aggregation of silver NPs, lowering their antibacterial activity [20]. Differently from Panacek *et al*., Dong *et al*. [21] showed no development of phenotypically silver-resistant bacteria (as evaluated by minimum inhibitory concentration, MIC, values) after 75 days of cultivation with daily transfers (involving about 500 bacterial generations) of *Pseudomonas putida* to low concentrations of Ag^+^ or AgNPs (10% of the MIC value), although they showed the development of mutations that differed between two exposures. Therefore, the development of resistance to silver nanomaterials can also be phenotypic and material-specific, necessitating further investigation when developing new antimicrobial nanomaterials. It is also reported that exposure to metallic antimicrobials may lead to resistance towards conventional antibiotics due to the co-location of the respective resistance genes in the bacterial genome or having the same mechanism for conferring resistance, an efflux pump, for example [22].

However, since biocidal metal-based nanoparticles, e.g. AgNPs, exhibit multiple attack mechanisms towards microorganisms, such as ROS production, membrane, protein and DNA damage [23], it has been proposed that NPs are less likely than conventional antibiotics to induce resistance development [12,24,25]. This fact makes them promising candidates as novel antimicrobials. Combining/stabilizing biocidal NPs, e.g. AgNPs, with an antimicrobial (bio)polymer, e.g. chitosan, may enhance NPs antibacterial potency. The polysaccharide chitosan has been combined with metal-based nanomaterials (e.g. Ag, Cu, Au, ZnO, Fe_3_O_4_, TiO_2_) [26,27] to form (nano)composites for different biomedical applications. Chitosan is a biocompatible and antimicrobial polymer (produced by the deacetylation of the naturally abundant chitin) [28] with immunomodulating properties that support wound healing [29]. The complex antimicrobial mechanism of chitosan involves an electrostatic attraction with microbes, disruption of cell membranes, and interference with cellular functions [30]. Thus, the silver-chitosan nanocomposites may target various aspects that are vitally important for microbes and could effectively prevent the development of resistance.

In this study, we evaluated the antibacterial efficacy and potential of silver-chitosan nanocomposites (nAgCSs) and comparatively AgNO_3_ to induce the development of AMR and cross-resistance to conventional antibiotics. The research topic is timely as these data are currently scarce. Indeed, a search made on May 08, 2024, in WoS using a combination of search terms “silver AND antibiotic resistance AND cross-resistance” as (Topic) yielded just 23 papers starting from 1999 up to 2023.

In this study, silver-chitosan nanocomposites with the silver-chitosan weight ratios of 1:1 (nAgCS-1) and 1:3 (nAgCS-3) were synthesised and studied as a potential material for antimicrobial wound dressings. Ionic silver (AgNO_3_) was evaluated as a reference for shedding Ag-ions from nAgCSs. Two clinical model bacterial strains were assessed: Gram-negative *Escherichia coli* ATCC 25922 and Gram-positive *Staphylococcus aureus* ATCC 6538. The bacterial strains were chosen as *E. coli* frequently has shown resistance to ampicillin and tetracycline [31,32], and *S. aureus*, particularly methicillin-resistant *S. aureus* (MRSA), has exhibited resistance to oxacillin [33] and, occasionally, to ciprofloxacin and erythromycin [34].

The experiment to assess AMR development spanned five weeks with daily exposures of bacteria to subinhibitory concentrations of nAgCSs and AgNO_3_ and weekly determination of minimum inhibitory concentrations (MIC) and minimum bactericidal concentrations (MBC). The 5-week duration of the experiment was chosen based on the data indicating that silver dressings are considered safe to use in burn units for up to 4 weeks, involving 8-10 dressing changes [35] and ISO 10993-1:2020 [36] where contact time up to 30 days is considered as prolonged exposure. To be even more prudent, we extended the exposure time to 5 weeks and applied daily re-inoculation and exposure of bacteria to the studied silver compounds.

The Kirby-Bauer disk diffusion test [37] was conducted to evaluate the effect of long-term continuous exposure to the studied silver compounds on developing bacterial cross-resistance to the essential conventional antibiotics used in treating bacterial infections, such as cefoxitin, ciprofloxacin, chloramphenicol, gentamicin, aztreonam, cefotaxime, ceftazidime, ampicillin, meropenem, tetracycline, clindamycin, erythromycin, oxacillin, vancomycin. Among the antibiotics evaluated in this study, gentamicin, meropenem, vancomycin, and ciprofloxacin have been classified by WHO as critically important antimicrobials for medicine [38].

## 2. Material and Methods

### 2.1 Synthesis and characterisation of silver-chitosan nanocomposites

All the chemicals used to synthesize the silver-chitosan nanocomposites (nAgCSs) were purchased from Sigma-Aldrich (Germany) and were of chemical purity ≥98%. nAgCSs were synthesized *via* the reduction of silver nitrate (AgNO_3_) with trisodium citrate (Na_3_C_6_H_5_O_7_; TSC) and stabilized by low molecular weight chitosan (50–190 kDa, Sigma-Aldrich). Chitosan was dissolved in 1% (vol/vol) acetic acid at the concentration of 10 g/L and allowed for complete dissolution at room temperature for at least 1–2 days before nAgCS synthesis.

For nAgCSs synthesis, AgNO_3_ (1 g Ag/L), TSC (10 g/L) and citric acid (10 g/L) solutions were prepared in deionized (DI) water (Millipore Milli-Q®, Merck Life Science BV, Belgium) and filtrated through a 0.2 µm pore-sized syringe filter (Sarstedt, Germany). Chitosan was diluted ten times in DI water (1 g/L) and filtered through a 0.45 µm filter (Sarstedt, Germany). Then, 10 mL of AgNO_3_ was added to 56 mL DI water and heated to boiling on a magnetic stirrer (Heidolph, Germany). Subsequently, 4 mL of TSC was added, and the mixture kept at ∼100°C, 250 rpm, for the next 10 minutes to facilitate the formation of silver nanoparticles. The suspension was cooled to room temperature, pH regulated to ∼4.8 by 1% citric acid, and chitosan added under stirring at 200 rpm. nAgCSs with silver chitosan weight ratios of 1:1 (nAgCS-1) and 1:3 (nAgCS-3) were synthesized. Next, the nAgCSs were ultrasonicated by a probe sonicator (Branson Digital Sonifier®, USA) for 10 seconds at 10% of maximum power (400 W) and dialysed against 2 L of DI water for 48 h (changing the DI water after 24 h) using dialysis tubes with cut-off 12 kDa (Viskase, USA). The concentration of Ag in nAgCS was measured by atomic absorption spectroscopy (AAS) (Analytic Jena, contrAA 800, Germany). Synthesized nAgCSs were stored at ∼4°C in the dark for six weeks. The shape and size of nAgCSs were analyzed using transmission electron microscopy in scanning mode (STEM) in a Titan Themis 200 (FEI) microscope, a service commissioned by Tartu University, Estonia. The primary sizes of nAgCs were determined by measuring their diameter in the respective STEM images using ImageJ software.

#### 2.1.1. Hydrodynamic size and ξ-potential measurements

Dynamic Light Scattering (DLS) and Electrophoretic Light Scattering (ELS) were used to characterize the nAgCSs. For the analysis, nAgCSs were suspended in DI water or 50% cation-adjusted Mueller-Hinton broth (CA-MHB_50_) (Oxoid Ltd, United Kingdom). Zetasizer Nano ZS and Zetasizer software 8.01 (Zetasizer Nano-ZS, Malvern Instruments Ltd, UK) were used for the analysis. Nanocomposites’ average hydrodynamic size (d_*H*_), surface charge (ζ-potential) and the suspension’s polydispersity index (PDI) were measured at nAgCSs concentration of 10 mg Ag/L in disposable cuvettes (DTS1061 folded capillary zeta cell or DTS0012 square polystyrene cuvette, Malvern Instruments Ltd, UK) at 25°C.

#### 2.1.2. UV-Vis analysis

Ultraviolet-visible (UV-Vis) absorption spectra of nAgCSs in DI water and CA-MHB_50_ were determined at 10 mg Ag/L in 2 mL quartz cuvettes (Hellma Analytics, Germany) at the wavelengths of 300–800 nm (step 2 nm) using a spectrophotometer (Multiscan Spectrum, Thermo Electron Corporation, Finland).

#### 2.1.3. Solubility/Dissolution

The concentration of silver ions shed from the nAgCSs in DI water and CA-MHB_50_ was measured at 10 mg Ag/L. A 12-well plate (Falcon) containing 3 mL of each diluted sample per well was incubated at 37°C (Biological thermostat BT 120) for 24 h to mimic the conditions of the bacterial susceptibility tests (cultivation medium composition, time, temperature). After 24 h, 2.7 mL of each sample was transferred into an Amicon Ultra-4 3K (3000 MWCO) filtration tube (Merck Millipore) and centrifuged for 40 minutes at 6225 g, 30°C (Sigma 3-16PK, Germany) to remove the non-soluble fraction of the nanocomposites. After centrifugation, 65% nitric acid (Honeywell), equivalent to 5% of the sample (filtrate) volume, was added, and the concentration of Ag ions measured by AAS (Analytic Jena, contrAA 800, Germany).

### 2.2. Analysis of antimicrobial efficiency and the development of antimicrobial resistance

#### 2.2.1. Bacterial strains and cultivation condition

Gram-negative bacteria *Escherichia coli* ATCC 25922 and Gram-positive bacteria *Staphylococcus aureus* ATCC 6538, obtained from the American Type Culture Collection (ATCC) were used. For the preparation of overnight culture, the bacteria were cultured in 3 ml of cation-adjusted Mueller-Hinton broth (CA-MHB) containing per 1 litre dehydrated infusion from beef 300 g, casein hydrolysate 17.5 g, starch 1.5 g, Ca^2+^ 22.5 mg and Mg^2+^ 11.25 mg at 37°C with shaking at 200 rpm.

#### 2.2.2. Antimicrobial efficiency evaluation

The antimicrobial efficiency of nAgCS-1, nAgCS-3, AgNO_3_ and low molecular weight chitosan (LMW CS) against bacteria *E. coli* ATCC 25922 and *S. aureus* ATCC 6538 was evaluated according to the antimicrobial susceptibility test protocol (ISO 20776-1:2019 [39]). In addition, a common antimicrobial, benzalkonium chloride (BAC, CAS no 63449-41-2, Sigma Aldrich, Germany), was used as a positive control. Briefly, an overnight bacterial culture was transferred to the fresh CA-MHB, adjusting the bacterial density to 5x10^5^ CFU/mL. Bacteria were incubated by mixing the bacterial suspension (75 μL) and with different concentrations of the studied silver compounds in DI water (75 μL) in 96-well transparent microplates (Falcon) at 37 °C for 24 h in the dark. The growth of bacteria was monitored by measuring the increase in absorbance at 600 nm in 30-minute intervals using a Spectramax Paradigm spectrophotometer (Molecular Devices, USA).

Based on the bacterial growth data, the following parameters were determined: i) minimum inhibitory concentration (MIC) – the lowest tested concentration of compounds where no visible growth of bacteria was observed and ii) effective concentrations (EC_20_ and EC_50_) of studied compounds – a concentration that reduced bacterial growth by 20 and 50%, respectively, compared to the control culture. The EC_20_ and EC_50_ values were derived from dose-response curves using the log-normal model with Microsoft Excel macro REGTOX [40]. Following the 24-hour incubation/exposure, 3 µl of each bacterial culture (control and those exposed to nAgCS, AgNO_3_ and BAC) was transferred to MHB agar plates and further incubated at 37°C for 24 h. After the incubation, colony formation was visually examined, and the minimum bactericidal concentration (MBC) was determined as the lowest tested concentration that completely inhibited bacterial growth (colony formation) on the agar plates.

#### 2.2.3. Antimicrobial resistance development test

In the AMR development experiments, bacteria were exposed to the studied silver compounds in CA-MHB_50_ for up to five weeks (35 days). In total, 35 successive 24-h sub-culture steps were conducted, involving repeated daily exposure to subinhibitory concentrations (EC_20_) of silver compounds: AgNO_3_ 0.8 and 1.6 mg Ag/L for *E. coli* and *S. aureus*, respectively, and nAgCS-1 6.0 mg Ag/L and nAgCS-3 3.0 mg Ag/L for both bacteria. In each sub-culture, bacteria were incubated in 3 mL of CA-MHB_50_ for 24 h in 14 mL round-bottom tubes (Falcon) with a subinhibitory concentration of silver compounds at 37°C, with shaking at 200 rpm (Fig. 1). In parallel, bacteria were also incubated in CA-MHB_50_ without any Ag compounds to serve as a non-treated (negative) control. The cultivation medium was renewed daily, transferring 3 µL of respective 24-h bacterial culture to 3 mL of fresh medium containing the studied Ag-compounds or not (negative control).

**Figure 1.**
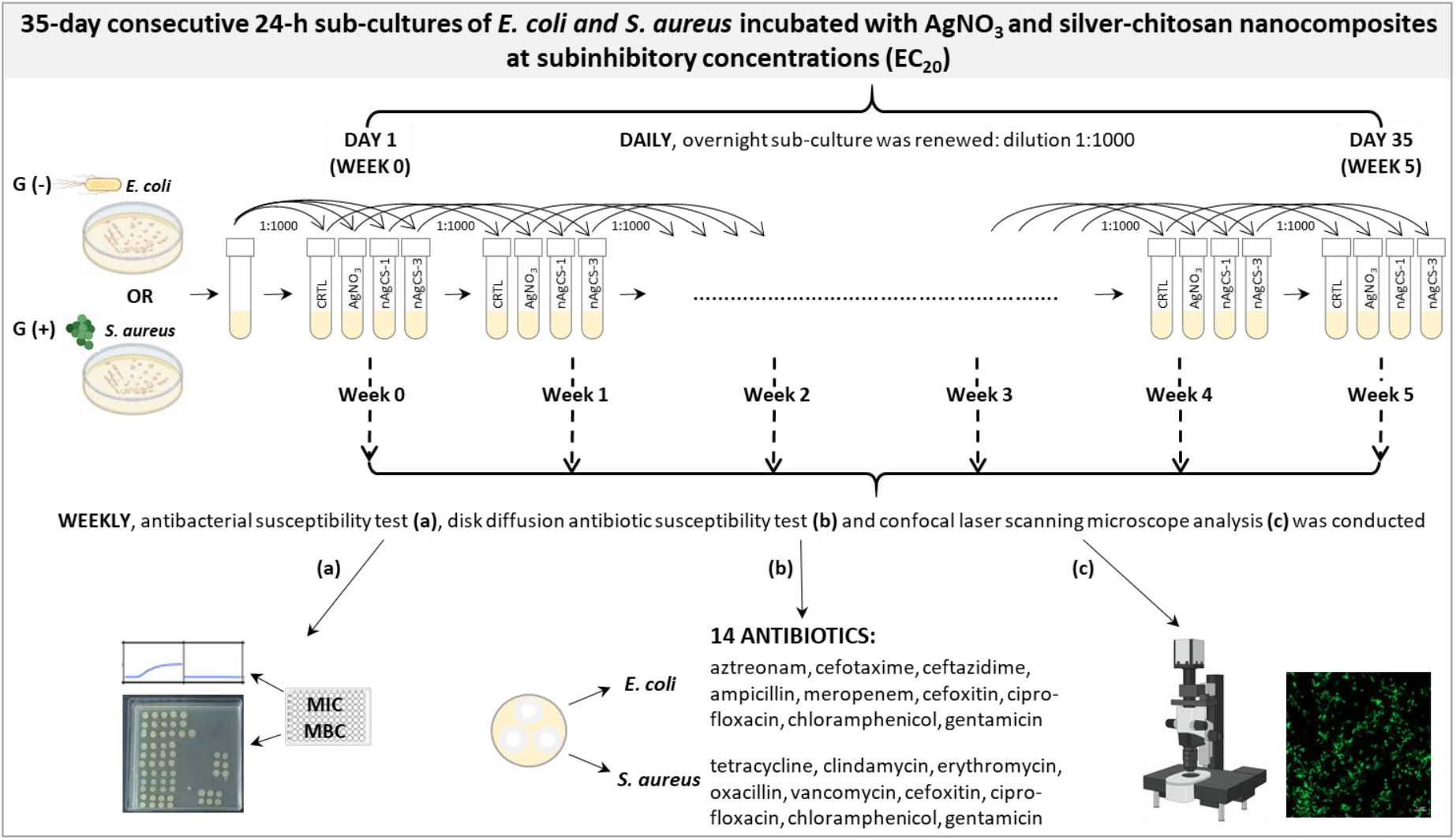
A scheme of the antimicrobial resistance development experiment.

Weekly, the bacteria’s susceptibility to (i) silver compounds and (ii) selected antibiotics was assessed using the antimicrobial susceptibility test, determining MIC and MBC values for nAgCSs and AgNO_3_ (see Section 2.2.2), and (ii) the disk diffusion susceptibility test determining the growth inhibition zones around the antibiotic disks (see Section 2.2.4), respectively. A methodology for assessing the development of AMR and cross-resistance was developed as part of this study.

#### 2.2.4. Disk diffusion susceptibility assay (Kirby-Bauer test)

To study the effect of continuous exposure to the silver compounds on the development of cross-resistance of bacteria to different antibiotics, the Kirby-Bauer disk diffusion susceptibility test [37] was performed. Bacteria exposed for 0–5 weeks to the studied silver compounds (Fig. 1) were weekly inoculated on the Mueller-Hinton agar and incubated for 20 h with the antimicrobial susceptibility disks containing 14 different antibiotics (Thermo Scientific, Oxoid; The European Committee on Antimicrobial Susceptibility Testing - EUCAST - disk content) (Tabel S1). The potential effect of continuous exposure to silver compounds on the development of cross-resistance of bacteria to different antibiotics was investigated by the formation of growth inhibition zones of bacteria around the antibiotics’ disks.

### 2.3. Confocal laser scanning microscopy

Bacteria exposed to the studied silver compounds for 0–5 weeks (Fig. 1) were analysed weekly by confocal laser scanning microscopy. Bacterial samples were diluted to the optical density (OD) at 600 nm of 1.0, and cells were stained for 15 min with 5 μM green fluorescent nucleic acid stain SYTO™ 9 (Invitrogen) at room temperature. Next, 20 µL of stained cell suspension was spread on the microscopy slide (Deltalab, Spain) and dried at 37 °C for 15 min. The samples were mounted in Mowiol (Sigma-Aldrich, Germany) and analysed with a confocal laser scanning microscope (CLSM) Zeiss LSM800 (Germany) equipped with a 100x oil immersion objective. To visualize Syto9-stained cells, the excitation/emission track settings of 488/505–550 nm were used. nAgCSs were visualized in reflection mode using a 640 nm laser. Images were processed with ZEN 2.6 software (Carl Zeiss Microscopy, Jena, Germany).

### 2.4. Statistical analysis

T-test analysis was conducted to identify statistical differences between determined values (MIC and MBC), and the results were displayed at a 95% confidence interval.

## 3. Results and discussion

### 3.1. Physico-chemical properties of the synthesised silver-chitosan nanocomposites

The silver-chitosan nanocomposites (100 mg Ag/L) with weight ratios of 1:1 (nAgCS-1) and 1:3 (nAgCS-3) were synthesized by reducing AgNO_3_ with trisodium citrate. According to STEM micrographs (Fig. S1) and ImageJ analysis, the primary sizes of nAgCS-1 and nAgCS-3 were 52 ± 18 and 47 ± 17 nm, respectively. The average hydrodynamic diameter (*d*_H_) of nAgCS-1 and nAgCS-3 in DI water was also in the nanoscale, measuring 91.8 and 78.8 nm, respectively (Table 1). However, in CA-MHB_50_, the hydrodynamic size of nAgCSs was in the range of 1-2 µm (Table 1), indicating the aggregation of nAgCS and/or protein corona formation in the growth medium [41]. The more significant change in *d*_H_ occurred in the nanocomposites with an Ag-chitosan weight ratio of 1:1 (nAgCS-1) compared to nAgCS-3 (the respective ratio was 1:3) in CA-MBH_50_, indicating the stabilizing role of chitosan in the nanocomposites.

**Table 1.** Physico-chemical properties of silver-chitosan nanocomposites (nAgCSs) in deionized (DI) water and 50% cation-adjusted Mueller-Hinton broth (CA-MHB_50_) at 10 mg Ag/L. The primary size of nAgCSs was ∼50 nm. The solubility of nAgCSs was determined at the concentration of 10 mg Ag/L after 24 h incubation in DI water and CA-MHB_50_ at 37°C.

| nAgCSs | Ag-chitosan<br>weight ratio | Hydrodynamic diameter <sup>a</sup>, nm<br>and (PDI)<sup>b</sup> |  | ζ-potential, mV |  | 24-h solubility, μg/L <sup>c</sup><br>(%) <sup>d</sup> |  |
| --- | --- | --- | --- | --- | --- | --- | --- |
|  |  | DI water | CA-MHB<sub>50</sub> | DI water | CA-MHB<sub>50</sub> | DI water | CA-MHB<sub>50</sub> |
| nAgCS-1 | 1:1 | 91.8±9.6<br>(0.35) | 2293±231<br>(0.29) | +30.8±2.2 | -4.16±0.3 | 156±52<br>(1.56%) | 2.03±1.3<br>(0.02%) |
| nAgCS-3 | 1:3 | 78.8±7.3<br>(0.46) | 1384±186<br>(0.50) | +32.5±2.0 | +3.12±0.2 | 25.1±3.3<br>(0.25%) | 3.07±1.7<br>(0.03%) |

<sup>a</sup>) intensity of scattered light diameter, <sup>b</sup>) PDI – polydispersity index, <sup>c</sup>) atomic absorption spectroscopy (AAS) analysis; <sup>d</sup>) - solubility %

The polydispersity (PDI) values of nAgCSs ranged from 0.29 to 0.50 in DI water and CA-MBH_50,_ suggesting moderate (PDI=0.1-0.4) to broad (PDI>0.4) polydisperse distribution [42]. The ζ-potential of nAgCSs in DI water was around +30 mV, but in CA-MHB_50_, these values shifted to -4.2 mV and +3.1 mV for nAgCS-1 and nAgCS-3, respectively (Table 1). Therefore, the suspension of nAgCSs in DI water, but not in CA-MHB_50_, could be considered theoretically stable (ζ-potential >30 [43]).

The UV-Vis analysis of nAgCSs showed maximum absorbance (*λ*_max_) at 418 nm in DI water and 422– 424 nm in CA-MBH_50_ (Fig. 2). The absorbance peak (SPR, surface plasmon resonance) at this range is typical of AgNPs [41]. In CA-MBH_50_, both nAgCSs SPR peaks shifted 4–6 nm to longer wavelengths (redshift), and the maximum absorbance value decreased compared to the absorption spectrum in DI water (Fig. 2), suggesting that nAgCSs may have aggregated.

**Figure 2.**
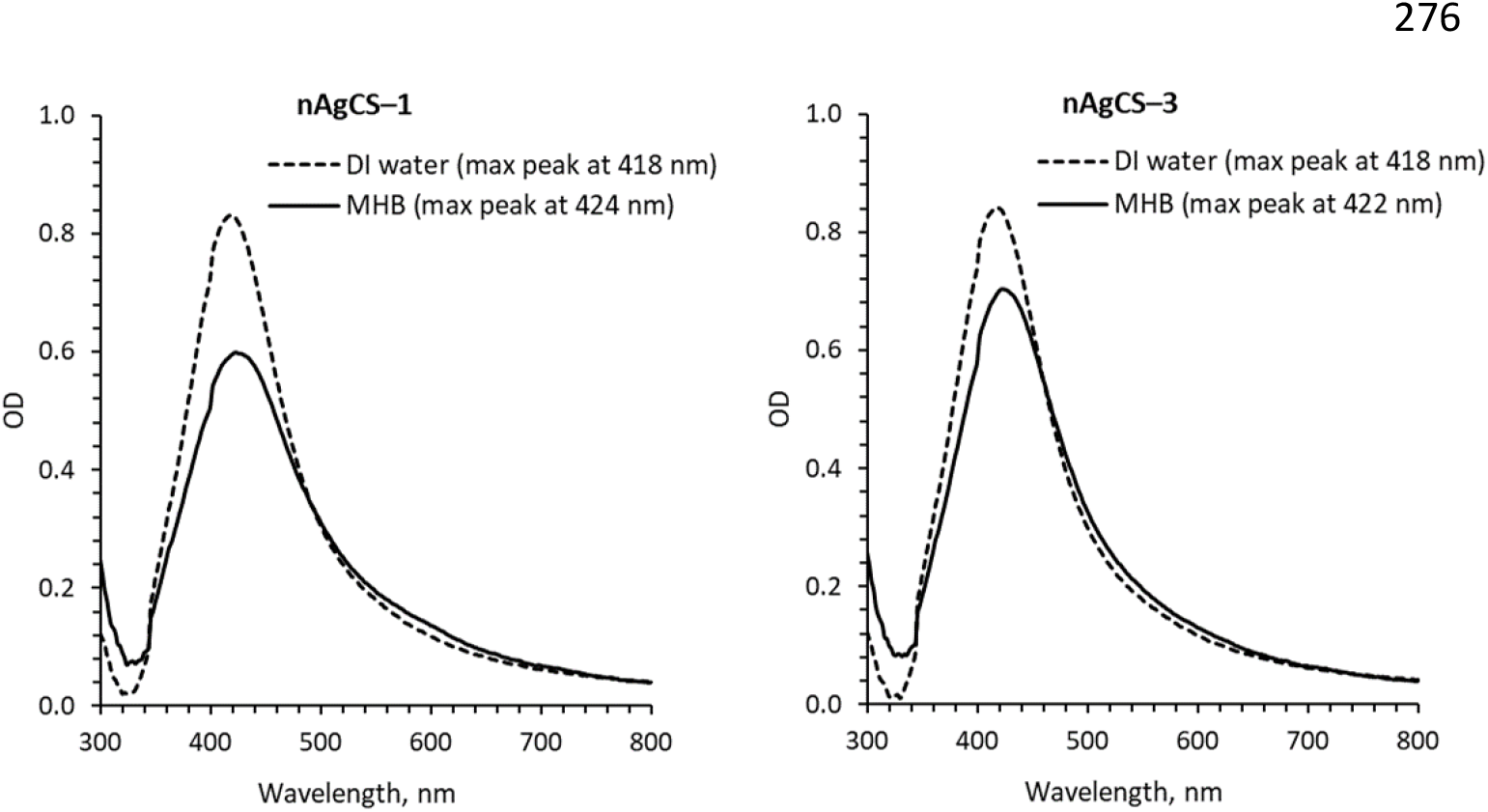
UV-Visible (UV-Vis) absorption spectra of silver-chitosan nanocomposites with Ag-chitosan weight ratios of 1:1 (nAgCS-1) and 1:3 (nAgCS-3) in deionized (DI) water and 50% cation-adjusted Mueller-Hinton broth (CA-MHB) at 10 mg Ag/L.

**Figure 2.**
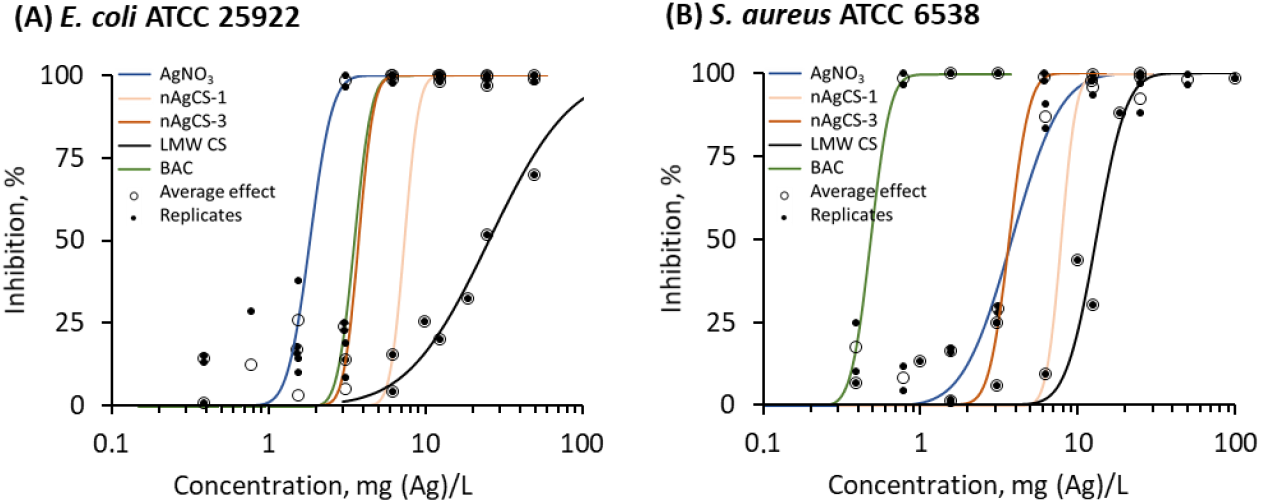
Inhibition of growth (% of control) of Gram-negative bacteria *Escherichia coli* (A) and Gram-positive bacteria *Staphylococcus aureus* (B) by AgNO_3_ (mg Ag/L), silver-chitosan nanocomposites (mg Ag/L) with weight ratios of 1:1 (nAgCS-1) and 1:3 (nAgCS-3), low molecular weight chitosan (LMW CS, mg/L) and benzalkonium chloride (BAC, mg/L) after 24-h growth in 50% Mueller-Hinton medium at 37°C compared to the non-exposed control (24-h concentration-effect curves). Please note the logarithmic x-axis. The respective EC_20_ and EC_50_ values are presented in Table 2.

The quantification of the shedding of silver ions from nAgCSs was performed in DI water and CA-MHB_50_ (toxicity testing medium) at the concentration of 10 mg Ag/L after 24 h incubation at 37°C in abiotic conditions (i.e. without cells). In DI water, the release of Ag ions from nAgCS-1 and nAgCS-3 was 1.6% and 0.3%, respectively (Table 1). nAgCS-3 was about 6-fold less soluble than nAgCS-1, indicating that higher chitosan concentration in nAgCS reduced the release of Ag ions. However, in CA-MHB_50_, the release of Ag ions from nAgCS-1 and nAgCS-3 was 9–27 fold lower than in DI water, comprising about 0.03% (Table 1).

### 3.2. Antimicrobial efficacy of silver compounds

The current study evaluated the development of AMR to nAgCSs and AgNO_3_ and cross-resistance to conventional antibiotics upon long-term exposure (up to 5 weeks) to subinhibitory levels of silver compounds. Before the AMR development experiments, the initial antibacterial efficacy (growth inhibition by 20% and 50%, MIC and MBC values) of nAgCSs, AgNO_3_, low molecular weight chitosan and benzalkonium chloride to bacteria *E. coli* ATCC 25922 and *S. aureus* ATCC 6538 was evaluated. The growth inhibition results (growth curves) are presented in the Supplementary Information (Fig. S2), and the respective dose-response curves in Fig. 2. Minimum inhibition concentration and (MIC) and minimum biocidal concentration (MBC) values are presented in Table 2.

**Table 2.** Effective concentrations causing 20% (EC_20_) and 50% (EC_50_) inhibition of growth, minimum inhibitory (MIC) and biocidal concentrations (MBC) of silver-chitosan nanocomposites with weight ratios of 1:1 (nAgCS-1) and 1:3 (nAgCS-3), AgNO_3_, low molecular chitosan (LMW CS) and benzalkonium chloride (BAC) to Gram-negative (G-) *Escherichia coli* ATCC 25922 and Gram-positive (G+) *Staphylococcus aureus* ATCC 3638 after 24 h growth in 50% Mueller-Hinton medium at 37°C. All concentrations are nominal (mg Ag/L or mg/L in the case of chitosan). EC_20_ and EC_50_ values are calculated from Fig. 2.

| Compounds | *Escherichia coli* ATCC 25922 (G-) |  |  |  | *Staphylococcus aureus* ATCC 6538 (G+) |  |  |  |
| --- | --- | --- | --- | --- | --- | --- | --- | --- |
|  | EC<sub>20</sub> | EC<sub>50</sub> | MIC | MBC | EC<sub>20</sub> | EC<sub>50</sub> | MIC | MBC |
| AgNO<sub>3</sub> | 1.49 | 1.85 | 3.13 | 3.13 | 2.37 | 3.81 | 7.29 | 12.5 |
| nAgCS-1 | 6.44 | 7.41 | 12.5 | 12.5 | 6.78 | 7.84 | 14.6 | 18.8 |
| nAgCS-3 | 3.27 | 3.78 | 6.25 | 6.25 | 3.01 | 3.67 | 6.25 | 6.25 |
| LMW CS | 11.5 | 25.9 | 70.0 | 100 | 10.5 | 13.2 | 25.0 | 37.5 |
| BAC | 3.00 | 3.60 | 6.25 | 8.59 | 0.40 | 0.48 | 0.91 | 3.13 |

It is well-known that AgNO_3_ (Ag-ions) is an efficient antibacterial agent acting at concentrations from 0.004 to 108 mg/L (median value of the MIC 3.3 mg/L; Bondarenko *et al*. [11]). In our study, AgNO_3_ inhibited bacterial growth at 3.1–7.3 mg/L after 24 h in CA-MHB_50_ at 37°C. Notably, Ag-ions were about 2–4 times more inhibitory to *E. coli* ATCC 25922 than to *S. aureus* ATCC 6538 (Fig. 2, Table 2), consistent with our earlier data [44,45]. On the contrary, chitosan and BAC were more toxic to *S. aureus* ATCC 6538 than *E. coli* ATCC 25922. The concentration-effect curve patterns of chitosan were also different, with a much steeper response curve in the case of *S. aureus* ATCC 6538 (Fig. 2). The MIC value of chitosan for *E. coli* ATCC 25922 and *S. aureus* ATCC 6538 was 70 and 25 mg/L, respectively (Table 2). Interestingly, the combination of silver and chitosan in the ratio of 1:3 yielded nanocomposites that were as efficient as AgNO_3_ against *S. aureus* ATCC 6538 (MIC 6.25 mg Ag/L) and as efficient as BAC against *E. coli* ATCC 25922 (MIC 6.25 mg/L). Thus, combining AgNPs and chitosan may result in nanocomposites with enhanced antibacterial efficacy. Generally, the silver-chitosan nanocomposites exhibited comparable antibacterial efficiency to both studied bacteria, with nAgCS-3 being about twice as efficient as nAgCS-1 (Table 2). The MBC values of AgNO_3_, nAgCS-1, nAgCS-3 and chitosan were comparable with MIC or ∼2-fold higher (Table 2), showing that all the studied compounds had bactericidal (not bacteriostatic) activity [46]. The review by Bondarenko *et al*. [11] of the antibacterial potency of silver nanoparticles demonstrated that MIC values ranged from 0.5-250 mg Ag/L, with a median MIC value of 7.1 mg Ag/L, showing that the MIC values obtained for the studied nAgCSs in the current study were in line with previously reported values for various bacteria and types of silver NPs.

Based on the antimicrobial efficacy results, subinhibitory concentrations (∼EC_20_) of silver compounds (0.8 and 1.6 mg Ag/L of AgNO_3_ for *E. coli* and *S. aureus*, respectively; 6.0 and 3.0 mg Ag/L of nAgCS-1 and nAgCS–3, respectively, for both bacteria) were selected for further AMR experiments to evaluate whether prolonged exposure (up to five weeks) may induce the bacteria to develop resistance to the compounds studied and/or cross-resistance to commonly used antibiotics.

### 3.3. Development of resistance to silver compounds

The overall AMR experiment lasted five consecutive weeks (35 days). During this period, 35 successive 24-h cultivation steps were performed daily with *E. coli* ATCC 25922 and *S. aureus* ATCC 6538, exposed to subinhibitory concentrations of AgNO_3_ and nAgCSs (Table 2). Weekly assessments of MBC and MIC for the silver compounds were conducted using an antibacterial susceptibility test. Simultaneously, control bacteria (not exposed) were sub-cultured consecutively for five weeks and exposed weekly to the studied silver compounds for the MIC and MBC determination (Fig. 1).

Although there was some statistically significant difference in MIC and/or MBC values of silver-exposed and control (not exposed) bacteria (Fig. 3, Table S2) during the 5-week exposure period, the resistance cut-off is defined as a greater than or equal to a 4-fold increase in the MIC/MBC [47] or even 8-fold [48]. Indeed, due to the exposure of 0–5 week *S. aureus* ATCC 6538 to nAgCS-3, a slight increase in MBC (but not in MIC) was registered (Table S2, Fig. 3). However, there were no statistically significant differences between the *S. aureus* nAgCS-3 MBC for exposed and non-exposed (control) groups, except for the 4-week MBC.

**Figure 3.**
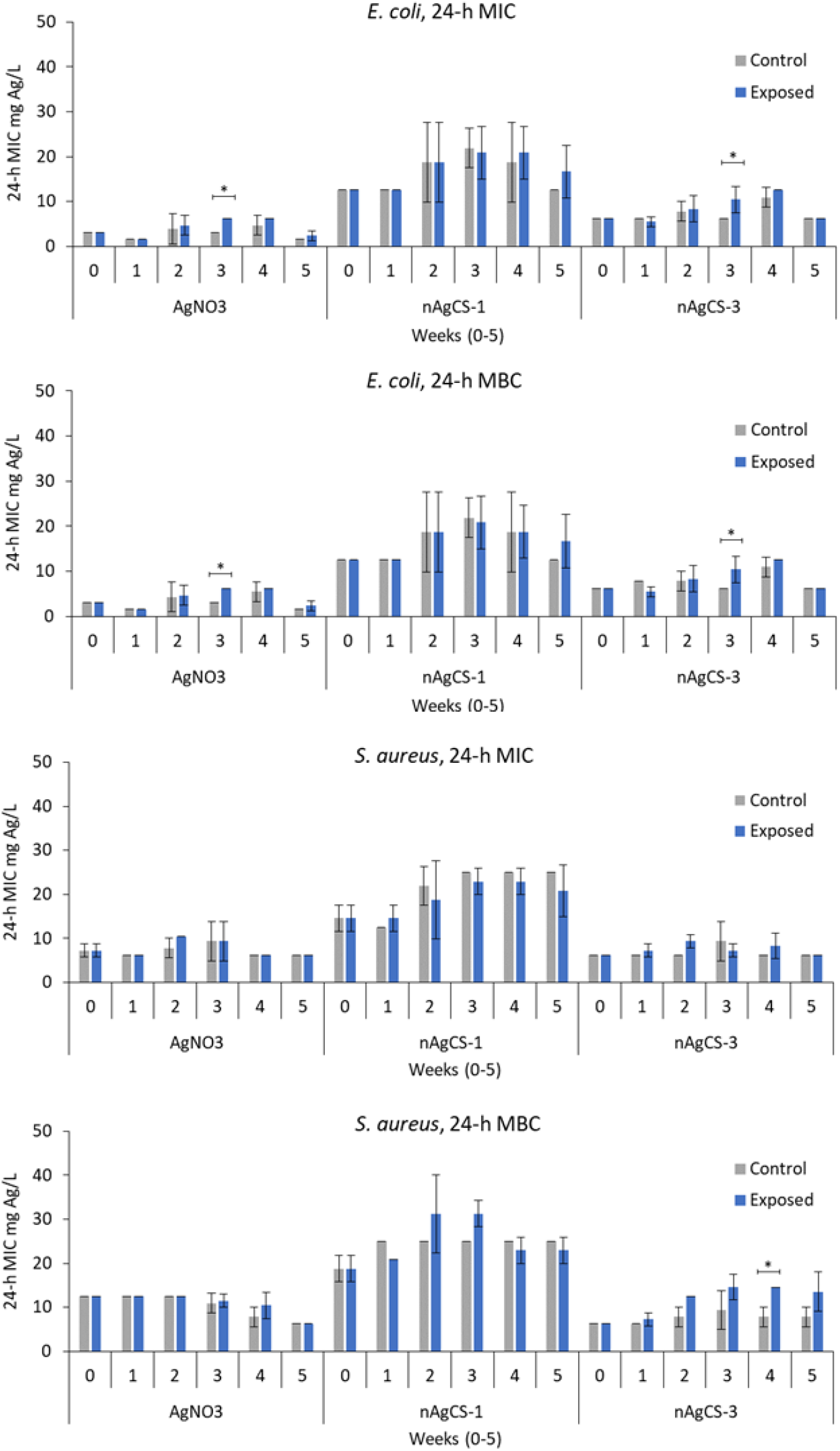
Minimum inhibitory concentration (MIC) and minimum biocidal concentration (MBC) of AgNO_3_ and silver-chitosan nanocomposites (nAgCS-1 and nAgCS-3) to *Escherichia coli* and *Staphylococcus aureus* after 24-h growth in CA-MHB50 at 37°C. The bacteria were continuously exposed to AgNO_3_ (0.8 mg Ag/L for *E. coli* and 1.6 mg Ag/L for *S. aureus*), nAgCS-1 (6.0 mg Ag/L) or nAgCS-3 (3.0 mg Ag/L) for 0–5 weeks in successive 24-hour culture steps. The control bacteria, not exposed to silver compounds, were sub-cultured under the same test conditions. MIC and MBC were determined weekly using the antibacterial susceptibility test (ISO 20776-1:2019). Asterisks designate to statistical differences (p ≤0.05).

Furthermore, the 0 and 5-week MBC values of nAgCS-3 for *S. aureus* were 6.25 and 13.5 mg Ag/L, respectively. Thus, the difference was approximately two-fold, which is lower than the criterion for resistance development. Consequently, the obtained results demonstrated that *E. coli* ATCC 25922 and *S. aureus* ATCC 6538 did not develop resistance to the tested silver compounds after continuous five weeks of exposure to AgNO_3_ and silver-chitosan nanocomposites at subinhibitory concentrations.

In contrast to our study, Panáček *et al*. [20] showed that the consecutive exposure of *E. coli* CCM 3954 to the subinhibitory concentration of colloidal silver induced the development of AMR: the MIC increased from 3.38 to 108 mg/L (32-fold) after 20 consecutive culture steps. The developed resistance was phenotypic and was triggered by the production of flagellin – an extracellular bacterial protein forming motility-enabling flagellar filaments – that caused the aggregation of silver NPs, lowering their antibacterial activity [20]. However, similar to our findings, Panacek *et al*. [20] did not observe the development of resistance to AgNO_3_. Thus, the development of antimicrobial resistance to nanosilver appears to depend on the type of nanoparticles used.

### 3.4. Development of cross-resistance to conventional antibiotics

In addition to investigating the development of AMR to AgNO_3_ and nAgCSs, the potential of these silver compounds to induce cross-resistance was assessed. The AB inhibition zone diameters (mm) obtained from the disk diffusion test agreed with the EUCAST data [49]. The results showed that no development of cross-resistance to the selected antibiotics was observed for *E. coli* ATCC 25922 and *S. aureus* ATCC 6538 upon five weeks of exposure to subinhibitory concentrations of the studied silver compounds (Fig. 4, Fig. S3 and Table S3).

**Figure 4.**
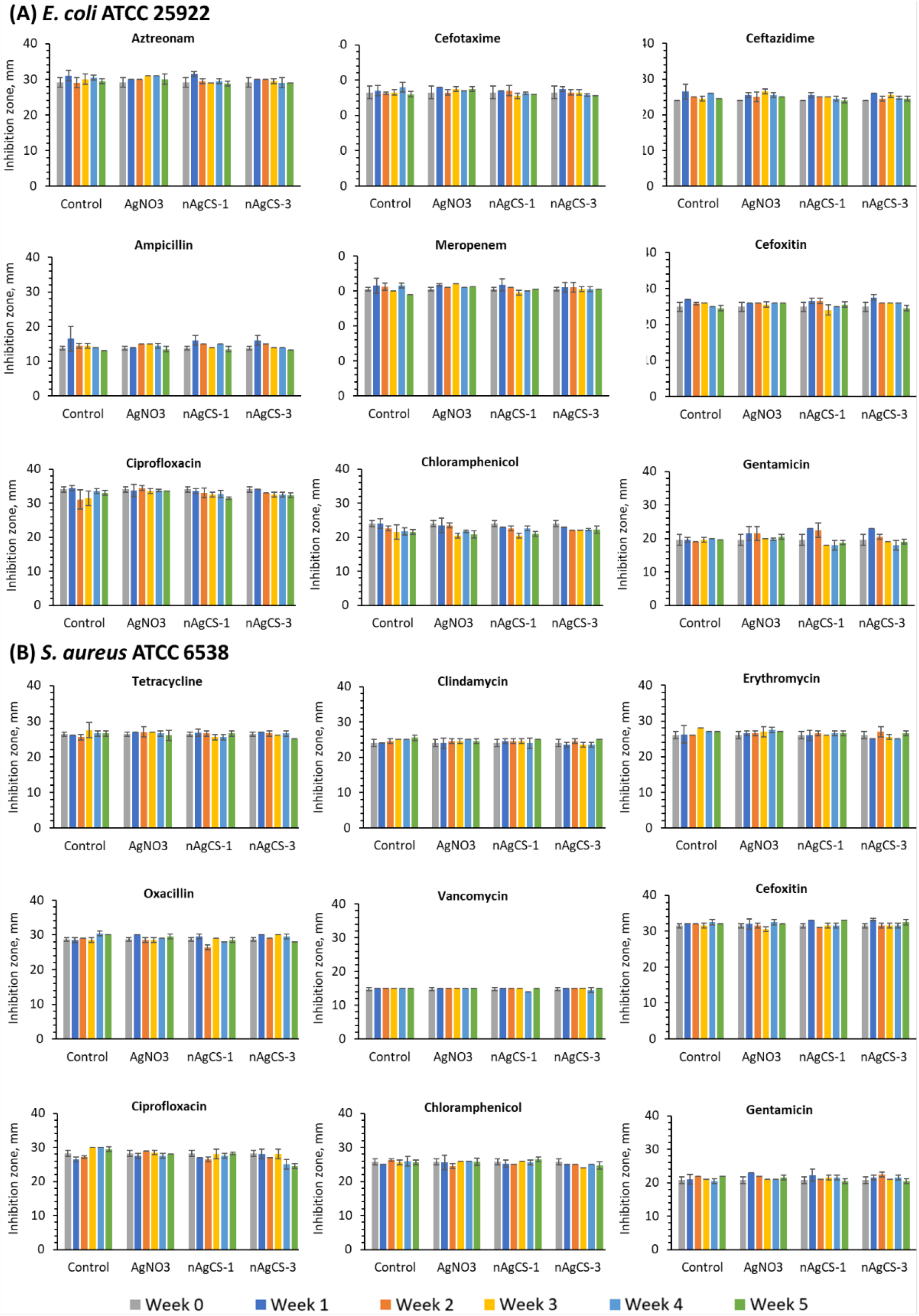
Susceptibility (formation of growth-inhibition zone, mm) of *Escherichia coli* (A) and *Staphylococcus aureus* (B) after 0–5 weeks of continuous exposure (or not; control) to AgNO_3_ and Ag-chitosan nanocomposites with weight ratios of 1:1 (nAgCS-1) and 1:3 (nAgCS-3) to the antibiotics (µg/disk): aztreonam (30), cefotaxime (5), ceftazidime (10), ampicillin (10), meropenem (10), cefoxitin (30), ciprofloxacin (5), chloramphenicol (30), gentamicin (10), tetracycline (30), clindamycin (2), erythromycin (15), oxacillin (1) and vancomycin (5).

### 3.5. Confocal laser scanning microscopy

Confocal laser scanning microscopy (CLSM) analyses were performed to assess the nAgCSs bacterial interactions (adsorption onto the surface of bacteria) and their effect on the morphology (e.g. clamping of the cells). Previous studies have shown that positively charged AgNPs may interact with microbial cells (adsorb onto the surface of cells). The particle-cell interactions increase the bioavailability and toxicity of silver to microorganisms, most probably via increased shedding of silver ions at the microbe-nanoparticle contact phase [11,50].

Interestingly, we observed that nAgCSs, especially nAgCS-3, promoted the formation of bacterial clusters (Fig. 5-6). Significantly pronounced clusters of *S. aureus* ATCC 6538 were observed already after the first week of exposure to nAgCS-3. In the case of *E. coli* ATCC 25922, clusters appeared later (in three weeks) and were not as massive, highlighting a specificity of nAgCSs towards Gram-positive bacteria. However, nAgCS-3 was almost equally toxic to both bacteria. This cluster-forming ability could explain why nAgCS-3 was twice as toxic as nAgCS-1 against the studied bacteria (Table 2).

**Figure 5.**
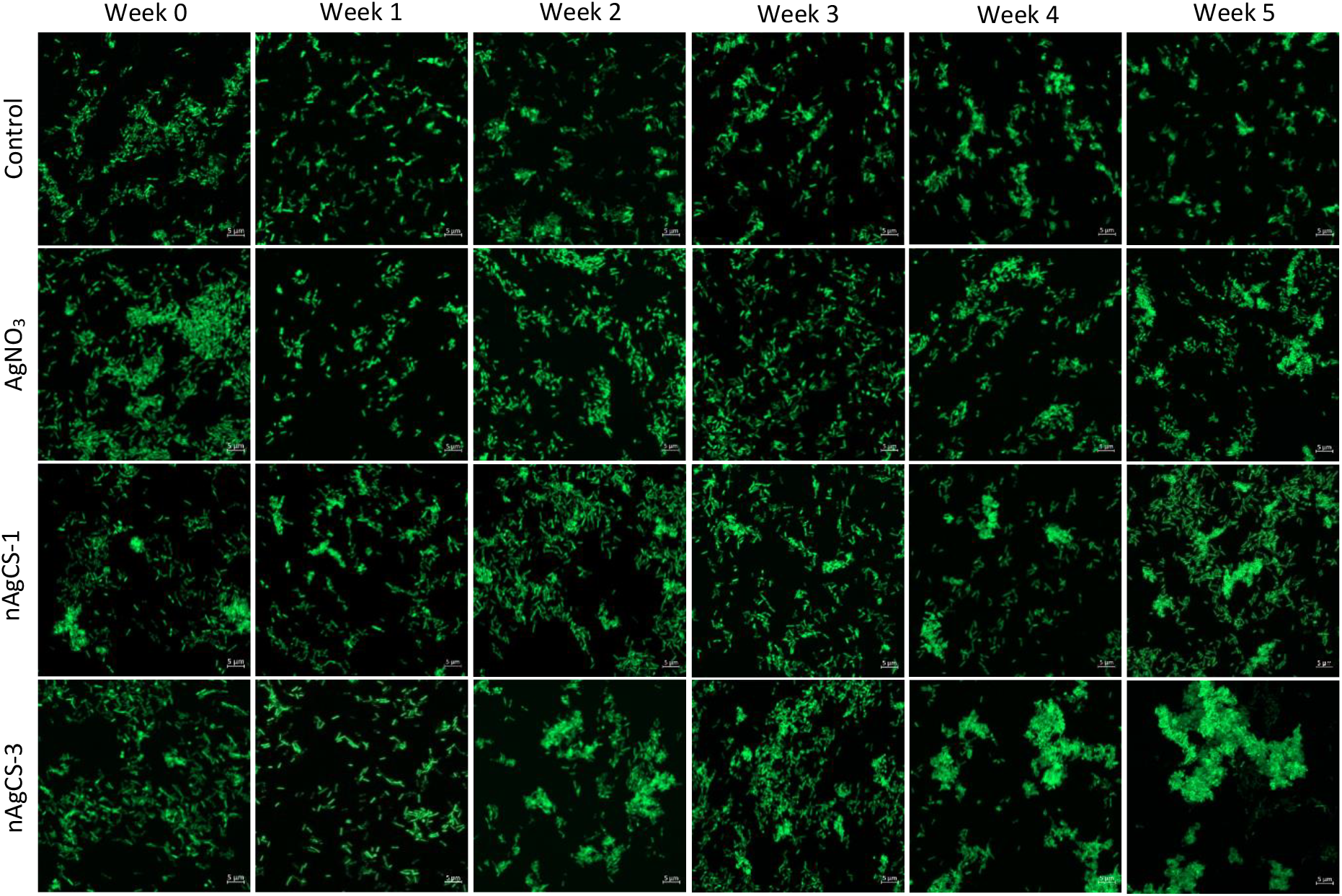
Confocal laser scanning microscopy images of *Escherichia coli*. The bacteria had been exposed (or not; control culture) to AgNO_3_ (0.8 mg Ag/L) and silver-chitosan nanocomposites (nAgCSs) with weight ratios of 1:1 (nAgCS-1, 6.0 mg Ag/L) or 1:3 (nAgCS-3, 3.0 mg Ag/L) for 0–5 weeks in consecutive 24-hour sub-cultures in 50% Mueller-Hinton medium at 37°C. Bacteria were stained by the green-fluorescent stain Syto-9 (nucleic acid probe), and nAgCSs were visualized in laser (640 nm) reflection mode (nAgCSs were not detected). All the scale bars are 5 µm.

Assumingly, the chitosan moiety brings chitosan-silver nanocomposites close to bacteria, allowing intimate contact between the bacterial cell surface and antimicrobial materials – chitosan and silver – to enhance their antibacterial action. Moreover, it has been demonstrated that NPs may induce cell aggregation due to cell membrane disruption or through the induction of stress response and the production of extracellular polymeric substances (EPS), which leads to the assembly of cells [51].

The laser-reflection mode [52] in CLSM analysis was also employed to assess the interactions between nAgCSs and bacteria. However, only nAgCS-3 was detectable within *S. aureus* ATCC 6538 clusters after 2 and 3 weeks of exposure (Fig. 6). This was not evident after 4–5 weeks, although the cell aggregation effect remained prominent. The fact that the studied nAgCSs tended to aggregate and settle during 24-hour incubation might explain why the nanocomposites were generally not detectable in the CLSM analysis.

**Figure 6.**
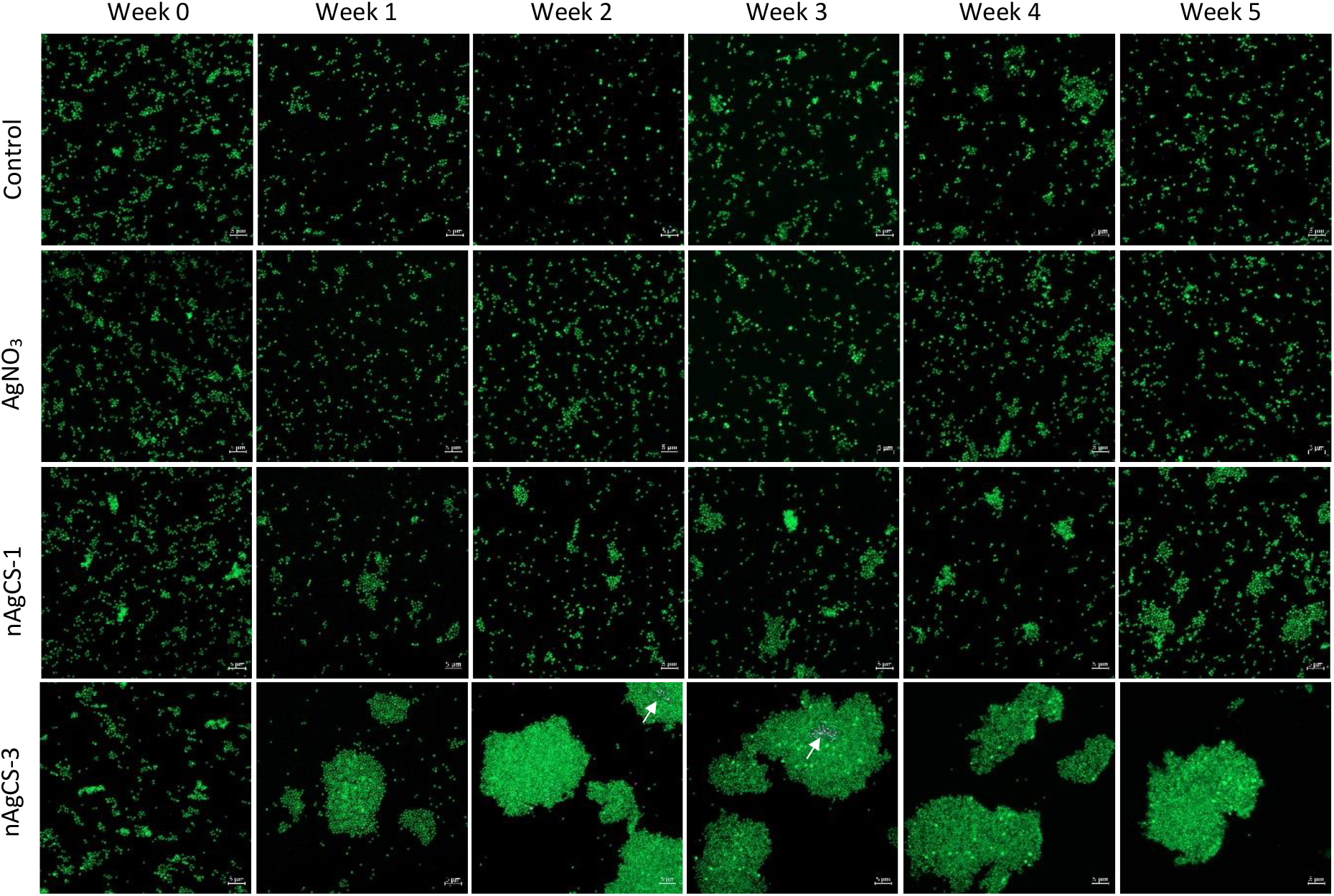
Confocal laser scanning microscopy images of *Staphylococcus aureus*. The bacteria had been exposed (or not in the case of control culture) to AgNO_3_ (1.6 mg Ag/L) and silver-chitosan nanocomposites (nAgCSs) with Ag-chitosan weight ratios of 1:1 (nAgCS-1, 6.0 mg Ag/L) or 1:3 (nAgCS-3, 3.0 mg Ag/L) for 0–5 weeks in consecutive 24-hour sub-cultures in Mueller-Hinton broth at 37°C. Bacteria were stained by the green-fluorescent stain Syto-9 (nucleic acid probe), and nAgCSs were visualized in laser (640 nm) reflection (purple points and white arrows, only seen in the images of nAgCS-3 after 2 and 3 weeks exposure). All the scale bars are 5 µm.

## 4. Conclusions

1. Silver-chitosan nanocomposites with a primary size of ∼50 nm and Ag-chitosan weight ratios of 1:1 (nAgCS-1) and 1:3 (nAgCS-3) were synthesized and characterised for physico-chemical properties.
2. The silver-chitosan nanocomposites showed antibacterial efficacy nearly equal to Ag-ions, acting at the minimum inhibitory concentration (MIC) level of 6–14 mg Ag/L against Gram-negative *E. coli* and Gram-positive *S. aureus*.
3. The development of phenotypical resistance (evaluated by MIC and MBC) in *E. coli* and *S. aureus* to either the nAgCSs or AgNO_3_ was not observed during the 5-week exposure to subinhibitory concentrations of the studied silver compounds. Importantly, also no cross-resistance towards 14 selected conventional antibiotics was developed in these bacteria.

Thus, silver-chitosan nanocomposites are promising candidates for antimicrobial applications, such as wound dressings and antimicrobial surfaces.

## Acknowledgements

This research was supported by the European Union and/or Estonian Research Council Grants PRG749, TEM-TA55, and the NAMUR+ core facility (TT 13).

## References

[1] United Nations Environment Programme, Bracing for superbugs: strengthening environmental action in the One Health response to antimicrobial resistance, Geneva, 2023. https://www.unep.org/resources/superbugs/environmental-action (accessed January 26, 2024).

[2] S. Ghosh, C. Bornman, M.M. Zafer, Antimicrobial Resistance Threats in the emerging COVID-19 pandemic: Where do we stand?, J Infect Public Health 14 (2021) 555–560. 10.1016/j.jiph.2021.02.011.

[3] World Health Organization, Antimicrobial resistance - Fact sheet, (2023). https://www.who.int/news-room/fact-sheets/detail/antimicrobial-resistance (accessed January 26, 2024).

[4] M. Xie, M. Gao, Y. Yun, M. Malmsten, V.M. Rotello, R. Zboril, O. Akhavan, A. Kraskouski, J. Amalraj, X. Cai, J. Lu, H. Zheng, R. Li, Antibacterial Nanomaterials: Mechanisms, Impacts on Antimicrobial Resistance and Design Principles, Angewandte Chemie International Edition 62 (2023). 10.1002/anie.202217345.

[5] M. Rai, A. Yadav, A. Gade, Silver nanoparticles as a new generation of antimicrobials, Biotechnol Adv 27 (2009) 76–83. 10.1016/j.biotechadv.2008.09.002.

[6] B. Nowack, H.F. Krug, M. Height, 120 Years of Nanosilver History: Implications for Policy Makers, Environ Sci Technol 45 (2011) 1177–1183. 10.1021/es103316q.

[7] N. Tripathi, M.K. Goshisht, Recent Advances and Mechanistic Insights into Antibacterial Activity, Antibiofilm Activity, and Cytotoxicity of Silver Nanoparticles, ACS Appl Bio Mater 5 (2022) 1391–1463. 10.1021/acsabm.2c00014.

[8] J.Y. Cheon, S.J. Kim, Y.H. Rhee, O.H. Kwon, W.H. Park, Shape-dependent antimicrobial activities of silver nanoparticles, Int J Nanomedicine Volume 14 (2019) 2773–2780. 10.2147/IJN.S196472.

[9] S. Tang, J. Zheng, Antibacterial Activity of Silver Nanoparticles: Structural Effects, Adv Healthc Mater 7 (2018). 10.1002/adhm.201701503.

[10] A.-C. Burduşel, O. Gherasim, A.M. Grumezescu, L. Mogoantă, A. Ficai, E. Andronescu, Biomedical Applications of Silver Nanoparticles: An Up-to-Date Overview, Nanomaterials 8 (2018) 681. 10.3390/nano8090681.

[11] O. Bondarenko, A. Ivask, A. Käkinen, I. Kurvet, A. Kahru, Particle-Cell Contact Enhances Antibacterial Activity of Silver Nanoparticles, PLoS One 8 (2013) e64060. 10.1371/journal.pone.0064060.

[12] R.Y. Pelgrift, A.J. Friedman, Nanotechnology as a therapeutic tool to combat microbial resistance, Adv Drug Deliv Rev 65 (2013) 1803–1815. 10.1016/j.addr.2013.07.011.

[13] C. Jelenko, Silver Nitrate Resistant E. coli, Ann Surg 170 (1969) 296–299.

[14] G.L. Mchugh, R.C. Moellering, C.C. Hopkins, M.N. Swartz, Salmonella Typhimurium resistant to silver nitrate, chloramphenicol, and ampicillin, The Lancet 305 (1975) 235–240. 10.1016/S0140-6736(75)91138-1.

[15] N.A. Nimer, Nosocomial Infection and Antibiotic-Resistant Threat in the Middle East, Infect Drug Resist Volume 15 (2022) 631–639. 10.2147/IDR.S351755.

[16] M. Nickol, J. Ciric, S. Falcinelli, D. Chertow, J. Kindrachuk, Characterization of Host and Bacterial Contributions to Lung Barrier Dysfunction Following Co-infection with 2009 Pandemic Influenza and Methicillin Resistant Staphylococcus aureus, Viruses 11 (2019) 116. 10.3390/v11020116.

[17] A. Gupta, S. Silver, Molecular Genetics: Silver as a biocide: Will resistance become a problem?, Nat Biotechnol 16 (1998) 888–888. 10.1038/nbt1098-888.

[18] K. Mijnendonckx, N. Leys, J. Mahillon, S. Silver, R. Van Houdt, Antimicrobial silver: uses, toxicity and potential for resistance, BioMetals 26 (2013) 609–621. 10.1007/s10534-013-9645-z.

[19] S.L. Percival, P.G. Bowler, D. Russell, Bacterial resistance to silver in wound care, Journal of Hospital Infection 60 (2005) 1–7. 10.1016/j.jhin.2004.11.014.

[20] A. Panáček, L. Kvítek, M. Smékalová, R. Večeřová, M. Kolář, M. Röderová, F. Dyčka, M. Šebela, R. Prucek, O. Tomanec, R. Zbořil, Bacterial resistance to silver nanoparticles and how to overcome it, Nat Nanotechnol 13 (2018) 65–71. 10.1038/s41565-017-0013-y.

[21] F. Dong, A.C. Quevedo, X. Wang, E. Valsami-Jones, J. Kreft, Experimental evolution of Pseudomonas putida under silver ion versus nanoparticle stress, Environ Microbiol 24 (2022) 905–918. 10.1111/1462-2920.15854.

[22] C. Pal, K. Asiani, S. Arya, C. Rensing, D.J. Stekel, D.G.J. Larsson, J.L. Hobman, Metal Resistance and Its Association With Antibiotic Resistance, in: 2017: pp. 261–313. 10.1016/bs.ampbs.2017.02.001.

[23] L. Wang, C. Hu, L. Shao, The antimicrobial activity of nanoparticles: present situation and prospects for the future, Int J Nanomedicine Volume 12 (2017) 1227–1249. 10.2147/IJN.S121956.

[24] V. Pareek, R. Gupta, S. Devineau, S.K. Sivasankaran, A. Bhargava, Mohd.A. Khan, S. Srikumar, S. Fanning, J. Panwar, Does Silver in Different Forms Affect Bacterial Susceptibility and Resistance? A Mechanistic Perspective, ACS Appl Bio Mater 5 (2022) 801–817. 10.1021/acsabm.1c01179.

[25] N.-Y. Lee, W.-C. Ko, P.-R. Hsueh, Nanoparticles in the Treatment of Infections Caused by Multidrug-Resistant Organisms, Front Pharmacol 10 (2019). 10.3389/fphar.2019.01153.

[26] G. Kravanja, M. Primožič, Ž. Knez, M. Leitgeb, Chitosan-Based (Nano)Materials for Novel Biomedical Applications, Molecules 24 (2019) 1960. 10.3390/molecules24101960.

[27] S. Murugesan, T. Scheibel, Chitosan-based nanocomposites for medical applications, Journal of Polymer Science 59 (2021) 1610–1642. 10.1002/pol.20210251.

[28] W. Wang, C. Xue, X. Mao, Chitosan: Structural modification, biological activity and application, Int J Biol Macromol 164 (2020) 4532–4546. 10.1016/j.ijbiomac.2020.09.042.

[29] H.B.T. Moran, J.L. Turley, M. Andersson, E.C. Lavelle, Immunomodulatory properties of chitosan polymers, Biomaterials 184 (2018) 1–9. 10.1016/j.biomaterials.2018.08.054.

[30] R. Zăvoianu, M. Tudorache, V.I. Parvulescu, B. Cojocaru, O.D. Pavel, New MgFeAl-LDH Catalysts for Claisen–Schmidt Condensation, Molecules 27 (2022) 8391. 10.3390/molecules27238391.

[31] D.A. Tadesse, S. Zhao, E. Tong, S. Ayers, A. Singh, M.J. Bartholomew, P.F. McDermott, Antimicrobial Drug Resistance in Escherichia coli from Humans and Food Animals, United States, 1950–2002, Emerg Infect Dis 18 (2012) 741–749. 10.3201/eid1805.111153.

[32] C.-M. Gaşpar, L.T. Cziszter, C.F. Lăzărescu, I. Ţibru, M. Pentea, M. Butnariu, Antibiotic Resistance among Escherichia coli Isolates from Hospital Wastewater Compared to Community Wastewater, Water (Basel) 13 (2021) 3449. 10.3390/w13233449.

[33] T.J. Foster, Antibiotic resistance in Staphylococcus aureus. Current status and future prospects, FEMS Microbiol Rev 41 (2017) 430–449. 10.1093/femsre/fux007.

[34] D. Pignataro, F. Foglia, M.T. Della Rocca, C. Melardo, B. Santella, V. Folliero, S. Shinde, P.C. Pafundi, F.C. Sasso, M.R. Iovene, M. Galdiero, G. Boccia, G. Franci, E. Finamore, M. Galdiero, Methicillin-resistant Staphylococcus aureus: epidemiology and antimicrobial susceptibility experiences from the University Hospital ‘Luigi Vanvitelli’ of Naples, Pathog Glob Health 114 (2020) 451–456. 10.1080/20477724.2020.1827197.

[35] K. Cutting, R. White, M. Edmonds, The safety and efficacy of dressings with silver – addressing clinical concerns, Int Wound J 4 (2007) 177–184. 10.1111/j.1742-481X.2007.00338.x.

[36] ISO 10993-1:2020. Biological evaluation of medical devices-Part 1: Evaluation and testing within a risk management process (ISO 10993-1:2018, including corrected version 2018-10), 2020. https://www.evs.ee/en/evs-en-iso-10993-1-2020 (accessed June 4, 2024).

[37] J. Hudzicki, Kirby-Bauer Disk Diffusion Susceptibility Test Protocol, American Society for Microbiology (2009). https://www.atcc.org.

[38] World Health Organization, Critically important antimicrobials for human hedicine, 6th revision 2018. Ranking of medically important antimicrobials for risk management of antimicrobial resistance due to non-human use, Geneva, 2019. https://iris.who.int/bitstream/handle/10665/312266/9789241515528-eng.pdf?sequence=1&isAllowed=y (accessed January 26, 2024).

[39] ISO 20776–1:2019. Susceptibility testing of infectious agents and evaluation of performance of antimicrobial susceptibility test devices. Part 1: Broth micro-dilution reference method for testing the in vitro activity of antimicrobial agents against rapidly growing aerobic bacteria involved in infectious diseases, (2019). https://www.evs.ee/et/iso-20776-1-2019 (accessed January 26, 2024).

[40] E. Vindimian, REGTOX Macro Excel, (2020). http://www.normalesup.org/~vindimian/en_index.html (accessed January 26, 2024).

[41] R. Vazquez-Muñoz, N. Bogdanchikova, A. Huerta-Saquero, Beyond the Nanomaterials Approach: Influence of Culture Conditions on the Stability and Antimicrobial Activity of Silver Nanoparticles, ACS Omega 5 (2020) 28441–28451. 10.1021/acsomega.0c02007.

[42] U. Nobbmann, Polydispersity – what does it mean for DLS and chromatography?, Malvern Panalytical Ltd (2017). https://www.materials-talks.com/polydispersity-what-does-it-mean-for-dls-and-chromatography/ (accessed January 26, 2024).

[43] Malvern Instruments Limited, Zeta potential - An introduction in 30 minutes. Technical note., (2015). http://www.malvernpanalytical.com (accessed January 26, 2024).

[44] S. Suppi, K. Kasemets, A. Ivask, K. Künnis-Beres, M. Sihtmäe, I. Kurvet, V. Aruoja, A. Kahru, A novel method for comparison of biocidal properties of nanomaterials to bacteria, yeasts and algae, J Hazard Mater 286 (2015) 75–84. 10.1016/j.jhazmat.2014.12.027.

[45] A.-L. Kubo, I. Capjak, I.V. Vrček, O.M. Bondarenko, I. Kurvet, H. Vija, A. Ivask, K. Kasemets, A. Kahru, Antimicrobial potency of differently coated 10 and 50 nm silver nanoparticles against clinically relevant bacteria Escherichia coli and Staphylococcus aureus, Colloids Surf B Biointerfaces 170 (2018) 401–410. 10.1016/j.colsurfb.2018.06.027.

[46] J.L. Davis, Pharmacologic Principles, in: Equine Internal Medicine, Elsevier, 2018: pp. 79–137. 10.1016/B978-0-323-44329-6.00002-4.

[47] D.J. Farrell, M. Robbins, W. Rhys-Williams, W.G. Love, Investigation of the Potential for Mutational Resistance to XF-73, Retapamulin, Mupirocin, Fusidic Acid, Daptomycin, and Vancomycin in Methicillin-Resistant Staphylococcus aureus Isolates during a 55-Passage Study, Antimicrob Agents Chemother 55 (2011) 1177–1181. 10.1128/AAC.01285-10.

[48] Clinical and Laboratory Standards Institute, M.P. Weinstein, Methods for dilution antimicrobial susceptibility tests for bacteria that grow aerobically, 2012.

[49] E. Matuschek, D.F.J. Brown, G. Kahlmeter, Development of the EUCAST disk diffusion antimicrobial susceptibility testing method and its implementation in routine microbiology laboratories, Clinical Microbiology and Infection 20 (2014) O255–O266. 10.1111/1469-0691.12373.

[50] K. Kasemets, S. Käosaar, H. Vija, U. Fascio, P. Mantecca, Toxicity of differently sized and charged silver nanoparticles to yeast Saccharomyces cerevisiae BY4741: a nano-biointeraction perspective, Nanotoxicology 13 (2019) 1041–1059. 10.1080/17435390.2019.1621401.

[51] B. Niu, G. Zhang, Effects of Different Nanoparticles on Microbes, Microorganisms 11 (2023) 542. 10.3390/microorganisms11030542.

[52] S. Käosaar, A. Kahru, P. Mantecca, K. Kasemets, Profiling of the toxicity mechanisms of coated and uncoated silver nanoparticles to yeast Saccharomyces cerevisiae BY4741 using a set of its 9 single-gene deletion mutants defective in oxidative stress response, cell wall or membrane integrity and endocytosis, Toxicology in Vitro 35 (2016) 149–162. 10.1016/j.tiv.2016.05.018.

