## Supplementary Material for "Do silver-chitosan nanocomposites promote bacterial resistance to silver or common antibiotics?"

### CONTENT

**Table S1.** List of antibiotics tested in disk diffusion susceptibility assay to analyse cross-resistance development of *Escherichia coli* and *Staphylococcus aureus* upon long-term exposure (0–5 weeks) to the subinhibitory level of the studied silver compounds (nAgCSs and $AgNO_3$). Antibiotic disks (Oxoid Ltd., United Kingdom). ........ 2

**Table S2.** Minimum inhibitory concentration (MIC, mg Ag/L) and minimum biocidal concentration (MBC, mg Ag/L) of $AgNO_3$ and silver-chitosan nanocomposites (nAgCS-1 and nAgCS-3) to *Escherichia coli* ATCC 25922 (A) and *Staphylococcus aureus* ATCC 6538 (B) after 24-h growth in 50% CA-MHB at 37°C. The bacteria were continuously exposed (E) (except control culture (C), not exposed to Ag compounds) to $AgNO_3$ (0.8 mg Ag/L for *E. coli* and 1.6 mg Ag/L for *S. aureus*), nAgCS-1 (6.0 mg Ag/L) or nAgCS-3 (3.0 mg Ag/L) for 0–5 weeks in successive 24-hour culture steps. MIC and MBC were determined weekly. ........ 3

**Table S3.** Inhibition zone diameters (mm; average ± Stdev) of the selected antibiotics from the disk diffusion test (Kirby–Bauer assay) observed for *E. coli* ATCC 25922 and *S. aureus* ATCC 6538 upon five-week exposure to sub-inhibitory concentrations of studied silver compounds ($AgNO_3$ 0.8 and 1.6 mg Ag/L for *E. coli* and *S. aureus*, respectively; nAgCS–1 6.0 and nAgCS–3 3.0 mg Ag/L for both bacteria). ........ 4

**Figure S1.** Scanning transmission electron microscopy (STEM) images of silver-chitosan nanocomposites (nAgCSs) with Ag-chitosan weight ratios of 1:1 (nAgCS-1, **A**) and 1:3 (nAgCS-3, **B**). .. 4

**Figure S2.** Growth of *Escherichia coli* ATCC 25922 (A) and *Staphylococcus aureus* ATCC 6538 (B) upon exposure to different concentrations of $AgNO_3$, silver-chitosan nanocomposites with Ag-chitosan weight ratios of 1:1 (nAgCS-1) and 1:3 (nAgCS-3), low molecular weight chitosan (LMW CS) and benzalkonium chloride (BAC) in 50% CA-MHB medium for 24 at 37°C. ........ 5

**Figure S3.** Representatives of disk diffusion agar plates from Kirby–Bauer test. White disks (Oxoid) containing antibiotics on an agar plate inoculated with *Staphylococcus aureus* ATCC 6538 upon five-week exposure to the sub-inhibitory concentrations of $AgNO_3$, nAgCS-1 and nAgCS-3. A zone of inhibition of the growth of the bacteria around the disk corresponds to the strain's susceptibility to the antibiotics (code description in Table S3). ........ 6

**Table S1.** List of antibiotics tested in disk diffusion susceptibility assay to analyse cross-resistance development of *Escherichia coli* and *Staphylococcus aureus* upon long-term exposure (0–5 weeks) to the subinhibitory level of the studied silver compounds (nAgCSs and $AgNO_3$). Antibiotic disks (Oxoid Ltd., United Kingdom).

| No | Antibiotic class | Antibiotic | Antibiotic code | Disk content, µg/disk | Disk diffusion susceptibility test | |
|---|---|---|---|---|---|---|
| | | | | | ***E. coli*** | ***S. aureus*** |
| 1 | Chloramphenicol | Chloramphenicol | C | 30 | X | X |
| 2 | Fluoroquinolones | Ciprofloxacin | CIP | 5 | X | X |
| 3 | Aminoglycosides | Gentamicin | CN | 10 | X | X |
| 4 | Cephalosporines | Cefoxitin | FOX | 30 | X | X |
| 5 | Cephalosporines | Cefotaxime | CTX | 5 | X | nd |
| 6 | Cephalosporines | Ceftazidime | CAZ | 10 | X | nd |
| 7 | Carbapenems | Meropenem | MEM | 10 | X | nd |
| 8 | Monobactams | Aztreonam | ATM | 30 | X | nd |
| 9 | Penicillins | Ampicillin | AMP | 10 | X | nd |
| 10 | Penicillins | Oxacillin | OX | 1 | nd | X |
| 11 | Macrolides | Erythromycin | E | 15 | nd | X |
| 12 | Glycopeptides | Vancomycin | VA | 5 | nd | X |
| 13 | Tetracyclines | Tetracycline | TE | 30 | nd | X |
| 14 | Lincosamides | Clindamycin | DA | 2 | nd | X |

nd – not determined

**Table S2.** Minimum inhibitory concentration (MIC, mg Ag/L) and minimum biocidal concentration (MBC, mg Ag/L) of $AgNO_3$ and silver-chitosan nanocomposites (nAgCS-1 and nAgCS-3) to *Escherichia coli* ATCC 25922 (A) and *Staphylococcus aureus* ATCC 6538 (B) after 24-h growth in 50% CA-MHB at 37°C. The bacteria were continuously exposed (E) (except control culture (C), not exposed to Ag compounds) to $AgNO_3$ (0.8 mg Ag/L for *E. coli* and 1.6 mg Ag/L for *S. aureus*), nAgCS-1 (6.0 mg Ag/L) or nAgCS-3 (3.0 mg Ag/L) for 0–5 weeks in successive 24-hour culture steps. MIC and MBC were determined weekly.

| *Escherichia coli* ATCC 25922 | | | | | | |
|---|---|---|---|---|---|---|
| **Weeks** | **$AgNO_3$** | | **nAgCS-1** | | **nAgCS-3** | |
| | 24-h MIC (C/E) | 24-h MBC (C/E) | 24-h MIC (C/E) | 24-h MBC (C/E) | 24-h MIC (C/E) | 24-h MBC (C/E) |
| 0 | 3.13/3.13 | 3.13/3.13 | 12.5/12.5 | 12.5/12.5 | 6.25/6.25 | 6.25/6.25 |
| 1 | 1.56/1.56 | 1.56/1.56 | 12.5/12.5 | 12.5/12.5 | 6.25/5.47 | 7.81/5.47 |
| 2 | 3.91/4.69 | 4.30/4.69 | 18.8/18.8 | 18.8/18.8 | 7.81/8.33 | 7.81/8.33 |
| 3 | 3.13/6.25 | 3.13/6.25 | 21.9/20.8 | 21.9/20.8 | 6.25/10.4 | 6.25/10.4 |
| 4 | 4.69/6.25 | 5.47/6.25 | 18.8/20.8 | 18.8/18.8 | 10.9/12.5 | 10.9/12.5 |
| 5 | 1.56/2.34 | 1.56/2.34 | 12.5/16.7 | 12.5/16.7 | 6.25/6.25 | 6.25/6.25 |
| *Staphylococcus aureus* ATCC 6538 | | | | | | |
| 0 | 7.29/7.29 | 12.5/12.5 | 14.6/14.6 | 18.8/18.8 | 6.25/6.25 | 6.25/6.25 |
| 1 | 6.25/6.25 | 12.5/12.5 | 12.5/14.6 | 25.0/20.8 | 6.25/7.29 | 6.25/7.29 |
| 2 | 7.81/10.4 | 12.5/12.5 | 21.9/18.8 | 25.0/31.3 | 6.25/9.38 | 7.81/12.5 |
| 3 | 9.38/9.38 | 10.9/11.5 | 25.0/22.9 | 25.0/31.3 | 9.38/7.29 | 9.38/14.6 |
| 4 | 6.25/6.25 | 7.81/10.4 | 25.0/22.9 | 25.0/22.9 | 6.25/8.33 | 7.81/14.6 |
| 5 | 6.25/6.25 | 6.25/6.25 | 25.0/20.8 | 25.0/22.9 | 6.25/6.25 | 7.81/13.5 |

**Table S3.** Inhibition zone diameters (mm; average ± Stdev) of the selected antibiotics from the disk diffusion test (Kirby–Bauer assay) observed for *E. coli* ATCC 25922 and *S. aureus* ATCC 6538 upon five-week exposure to sub-inhibitory concentrations of studied silver compounds ($AgNO_3$ 0.8 and 1.6 mg Ag/L for *E. coli* and *S. aureus*, respectively; nAgCS–1 6.0 and nAgCS–3 3.0 mg Ag/L for both bacteria).

| ***E. coli* ATCC 25922** | | | | | |
|---|---|---|---|---|---|
| **No** | **Antibiotics** | **Control** | **$AgNO_3$** | **nAgCS-1** | **nAgCS-3** |
| 1 | Cefoxitin (FOX30) | 25.7±0.9 | 25.9±0.3 | 25.5±1.2 | 26.0±1.1 |
| 2 | Aztreonam (ATM30) | 30.0±1,2 | 30.4±0.5 | 29.7±1.1 | 29.5±0.7 |
| 3 | Cefotaxime (CTX5) | 26.8±1.0 | 27.3±0.7 | 26.4±0.9 | 26.4±0.9 |
| 4 | Ceftazidime (CAZ10) | 25.3±1.2 | 25.5±0.9 | 24.8±0.6 | 25.1±0.7 |
| 5 | Ampicillin (AMP10) | 14.5±1.7 | 14.4±0.7 | 14.7±1.1 | 14.5±1.1 |
| 6 | Meropenem (MEM10) | 30.7±1.3 | 31.4±0.5 | 30.6±1.1 | 30.7±0.8 |
| 7 | Ciprofloxacin (CIP5) | 32.7±1.8 | 33.8±0.8 | 32.7±1.0 | 32.9±0.8 |
| 8 | Chloramphenicol (C30) | 22.3±1.4 | 22.0±1.6 | 21.9±1.2 | 22.3±0.5 |
| 9 | Gentamicin (CN10) | 19.5±0.5 | 20.7±1.3 | 20.1±2.5 | 19.9±1.9 |
| ***S. aureus* ATCC 6538** | | | | | |
| **No** | **Antibiotics** | **Control** | **$AgNO_3$** | **nAgCS-1** | **nAgCS-3** |
| 1 | Cefoxitin (FOX 30) | 32.0±0.5 | 31.7±1.0 | 32.0±0.9 | 32.1±0.9 |
| 2 | Tetracycline (TE30) | 26.4±1.1 | 26.7±0.8 | 26.2±0.8 | 26.2±0.8 |
| 3 | Clindamycin (DA2) | 24.8±0.6 | 24.5±0.7 | 24.5±0.7 | 24.0±0.8 |
| 4 | Erythromycin (E15) | 26.9±1.1 | 26.9±0.7 | 26.3±0.7 | 25.8±1.0 |
| 5 | Oxacillin (OX1) | 29.3±1.0 | 29.1±0.7 | 28.3±1.2 | 29.3±0.8 |
| 6 | Vancomycin (VA5) | 15.0±0 | 15.0±0 | 14.8±0.4 | 14.9±0.3 |
| 7 | Ciprofloxacin (CIP5) | 28.7±1.6 | 28.1±0.7 | 27.5±0.9 | 26.5±1.8 |
| 8 | Chloramphenicol (C30) | 25.7±0.8 | 25.6±1.0 | 25.7±0.8 | 24.8±0.5 |
| 9 | Gentamicin (CN10) | 21.3±0.8 | 21.7±0.8 | 21.4±0.9 | 21.4±0.8 |

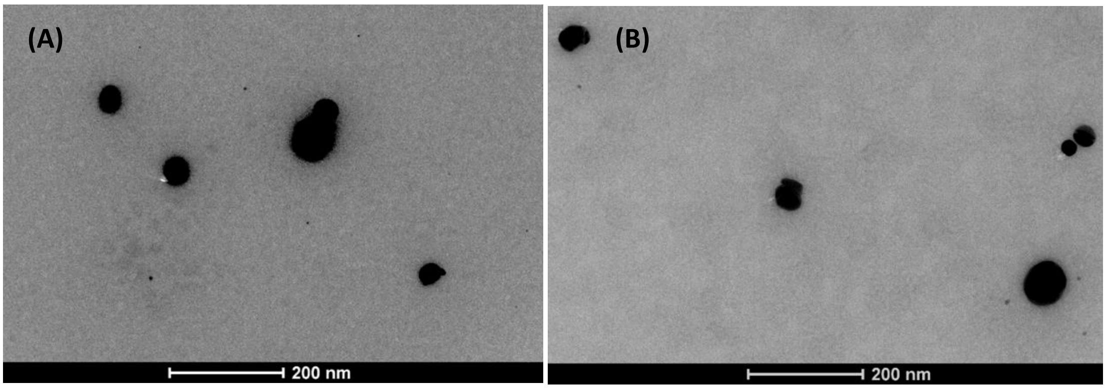


**Figure S1.** Scanning transmission electron microscopy (STEM) images of silver-chitosan nanocomposites (nAgCSs) with Ag-chitosan weight ratios of 1:1 (nAgCS-1, **A**) and 1:3 (nAgCS-3, **B**).

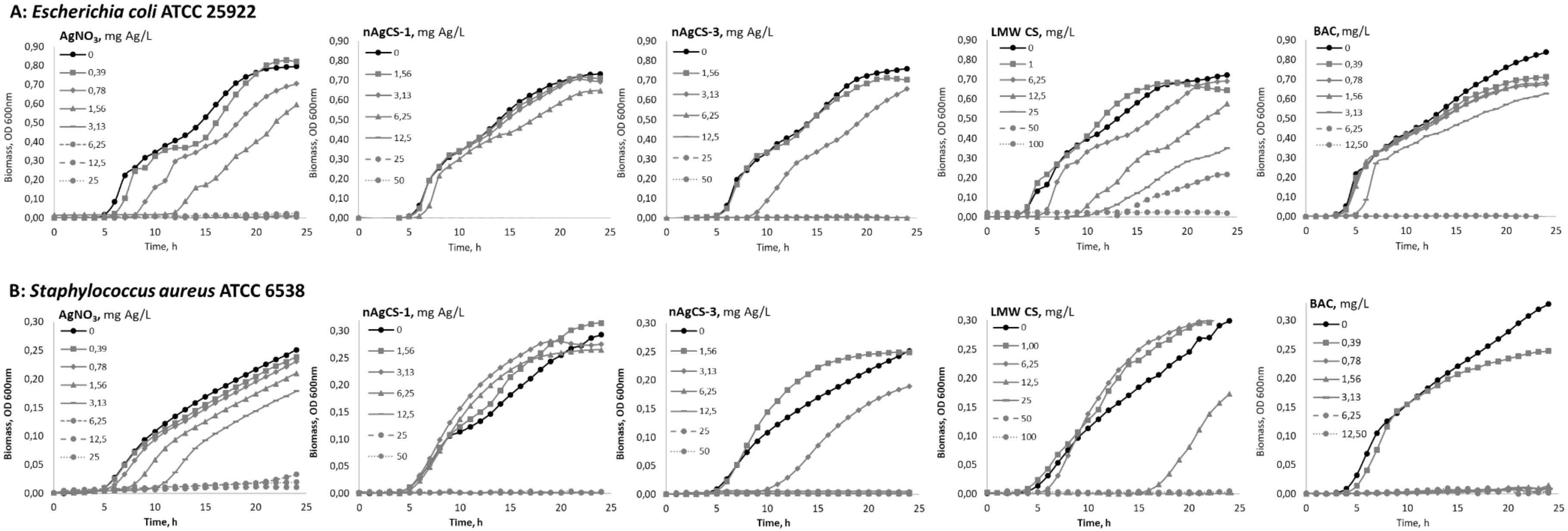


**Figure S2.** Growth of *Escherichia coli* ATCC 25922 (A) and *Staphylococcus aureus* ATCC 6538 (B) upon exposure to different concentrations of $AgNO_3$, silver-chitosan nanocomposites with Ag-chitosan weight ratios of 1:1 (nAgCS-1) and 1:3 (nAgCS-3), low molecular weight chitosan (LMW CS) and benzalkonium chloride (BAC) in 50% CA-MHB medium for 24 at 37°C.

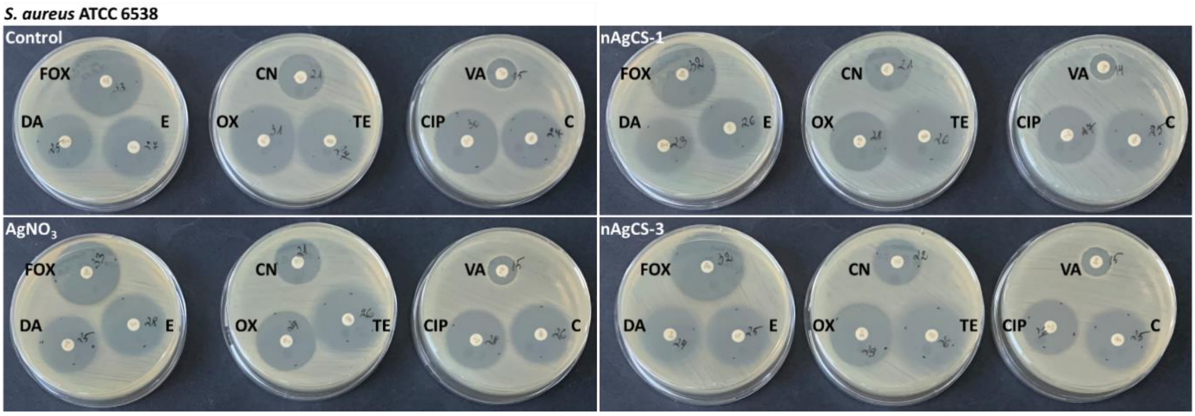


**Figure S3.** Representatives of disk diffusion agar plates from Kirby–Bauer test. White disks (Oxoid) containing antibiotics on an agar plate inoculated with *Staphylococcus aureus* ATCC 6538 upon five-week exposure to the sub-inhibitory concentrations of $AgNO_3$, nAgCS-1 and nAgCS-3. A zone of inhibition of the growth of the bacteria around the disk corresponds to the strain's susceptibility to the antibiotics (code description in Table S3).
